# The continuing impact of an ancient polyploidy on the genomes of teleosts

**DOI:** 10.1101/619205

**Authors:** Gavin C. Conant

**Affiliations:** Department of Biological Sciences, North Carolina State University, Raleigh, NC, U.S.A.; Bioinformatics Research Center, North Carolina State University, Raleigh, NC, U.S.A.; Program in Genetics, North Carolina State University, Raleigh, NC, U.S.A.; Division of Animal Sciences, University of Missouri, Columbia, MO, U.S.A.

**Keywords:** polyploidy, evolution of development, evolutionary model, sensory evolution, evolution of visual systems

## Abstract

The ancestor of most teleost fishes underwent a whole-genome duplication event three hundred million years ago. Despite its antiquity, the effects of this event are evident both in the structure of teleost genomes and in how those genes still operate to drive form and function. I describe the inference of a set of shared syntenic regions that survive from the teleost genome duplication (TGD) using eight teleost genomes and the outgroup gar genome (which lacks the TGD). I phylogenetically modeled the resolution of the TGD via shared and independent gene losses, concluding that it was likely an allopolyploidy event due to the biased pattern of these gene losses. Duplicate genes surviving from this duplication in zebrafish are less likely to function in early embryo development than are genes that have returned to single copy. As a result, surviving ohnologs function later in development, and the pattern of which tissues these ohnologs are expressed in and their functions lend support to recent suggestions that the TGD was the source of a morphological innovation in the structure of the teleost retina. Surviving duplicates also appear less likely to be essential than singletons, despite the fact that their single-copy orthologs in mouse are no less essential than other genes. Nonetheless, the surviving duplicates occupy central positions in the zebrafish metabolic network.

## Introduction

The study of doubled genomes (or polyploids) has a long history in genetics (Kuwada 1911; Clausen and Goodspeed 1925; Ohno 1970; Taylor and Raes 2004; Garsmeur, et al. 2013), but it was the advent of complete genome sequencing that most dramatically confirmed the role of polyploidy in shaping modern eukaryote genomes (Van de Peer, et al. 2017). The remnants of ancient genome duplications have been found across the eukaryotic phylogeny, from flowering plants (Soltis, et al. 2009) and yeasts (Wolfe and Shields 1997) to ciliates (Aury, et al. 2006) vertebrates (Ohno 1970; Kasahara 2007; Makino and McLysaght 2010) nematodes (Blanc-Mathieu, et al. 2017) and arachnids (Schwager, et al. 2017).

While flowering plants may be the “champions” of polyploidy (Soltis, et al. 2009), genome duplication has also extensively shaped the evolution of the teleost fishes (Chenuil, et al. 1999; Alves, et al. 2001; Braasch and Postlethwait 2012; Yang, et al. 2015). Polyploidies ranging in age from recent (<1Mya) hybridizations to very old events are known, including events shared among clades in the salmonids, carps, and sturgeons. The event considered here is a very old one that occurred between 320 and 400 Mya in the ancestor of most ray-fined fishes: the *teleost genome duplication* (TGD; Christoffels, et al. 2004; Hoegg, et al. 2004; Vandepoele, et al. 2004; Crow, et al. 2005; Sémon and Wolfe 2007). Evidence for this event started to accumulate in the late 1990s and became effectively irrefutable with the sequencing of the first teleost genomes (Aparicio, et al. 2002; Jaillon, et al. 2004; Van de Peer 2004).

Several evolutionary changes are associated with the TGD, including molecular divergence in vitamin receptors (Kollitz, et al. 2014), circulatory system genes (Moriyama, et al. 2016) and in the structure of core metabolism (Steinke, et al. 2006). Indeed, the classic example of duplicate gene divergence by subfunctionalization involves two zebrafish *ohnologs* (duplicates that are the products of a WGD; Wolfe 2000) *eng1a* and *eng1b* from the TGD (Force, et al. 1999). At the genome scale, the TGD probably increased the genome rearrangement rate for a period (Sémon and Wolfe 2007), as well as increasing the rate of sequence insertions and deletions in surviving ohnologs (Guo, et al. 2012). Likewise, extant teleost genomes show evidence for *reciprocal gene loss* between alternative copies of homologous genes created by the TGD, a pattern that can induce reproductive isolation between populations possessing it (Naruse, et al. 2004; Scannell, et al. 2006; Semon and Wolfe 2007).

A phylogenomic study of the TGD was undertaken by Inoue and colleagues (2015), who concluded that, as with other WGD events, the TGD was followed by an initial period of very rapid duplicate gene loss (Scannell, et al. 2007; McGrath, et al. 2014). However, their study was limited because it used gene tree/species tree reconciliation to identify the relics of the TGD, an approach which is known to have limitations relative to methods based on the identification and analysis of blocks of double-conserved synteny (DCS; Nakatani and McLysaght 2019; Zwaenepoel, et al. 2019). As a result, Inoue et al., could not completely phase the surviving genes in each genome relative to each other, making the process of estimating loss timings more challenging and suggesting that a new analysis of the TGD is appropriate. This opportunity is particularly important because zebrafish’s role as a developmental model gives us a unique opportunity to explore the effects of WGD on developmental evolution. That its effects may be important has long been hypothesized, with one example being the suspected role of Hox gene duplication in creating plasticity in body-plan evolution (Hoegg, et al. 2007). Moreover, much of the work on the “rules” of evolution after genome duplication have been performed using relatively recent events in flowering plants and yeasts, with less understanding of the very-long term effects of polyploidy. These proposed rules include the tendency of more highly interacting genes to remain as ohnolog pairs longer after WGD, a pattern explained by the *dosage balance hypothesis* (DBH; Freeling and Thomas 2006; Birchler and Veitia 2007; Edger and Pires 2009; Freeling 2009; Makino and McLysaght 2010; Conant, et al. 2014). The DBH argues that cellular interactions, governed as they are by biochemical kinetics (Veitia and Potier 2015), can be sensitive to imbalances in the concentrations of the interacting entities, driving those interacting genes to be maintained in similar dosages (e.g., as ohnologs after WGD). Importantly, the effects of the DBH may not preserve ohnologs indefinitely (Conant, et al. 2014), and the TGD is old enough to explore this question. A second rule of polyploidy is that the two parent genomes that contribute to an allopolyploid are not generally equal: instead *biased fractionation* is observed, whereby one of the two parent genomes retains more genes than does the other (Thomas, et al. 2006; Sankoff, et al. 2010; Schnable, et al. 2011; Tang, et al. 2012; Bird, et al. 2018; Emery, et al. 2018; Wendel, et al. 2018). The role of biased fractionation in the resolution of the TGD has also not, to my knowledge, been explored.

Using POInT, the <u>P</u>olyploid <u>O</u>rthology <u>In</u>ference <u>T</u>ool (Conant and Wolfe 2008), I analyzed the resolution of the TGD, based on nearly 5600 duplicated loci that are detectable in eight teleost genomes. I find that the surviving ohnologs produced by the TGD are distinct in their character even after more than 300 million years of evolution. They are rarely expressed in the earliest phases of development, are less likely to be essential in zebrafish, and they occupy more central positions in the zebrafish metabolic network. In addition, there are suggestions that the TGD helped shape a key innovation in the teleost visual system.

## Results

### Identifying the relics of the TGD in eight teleost genomes

Using a pipeline that places homologs of genes from gar (which lacks the TGD) into double-conserved synteny (DCS) blocks, I identified 5589 ancestral loci in eight teleost genomes surviving from the TGD (Figure 1). Each such locus was duplicated in the TGD and now has either one or both surviving genes from that event in each of the 8 genomes (c.f., Figure 1). I analyzed these blocks with POInT (Conant and Wolfe 2008), our tool for modeling the evolution of polyploid genomes. Because POInT infers orthologous chromosome segments based on a common gene order and shared gene losses along a phylogeny, it requires an estimate of the order of these loci in the ancestral genome immediately prior to the TGD (e.g., as was previously done for yeast; Gordon, et al. 2009). The TGD is considerably older and the genomes involved more rearranged than was the case for the yeast, *Arabidopsis* and grass polyploidies we previously analyzed (Conant and Wolfe 2008; Conant 2014; Emery, et al. 2018). Hence, I explored several means for estimating this order (*Methods*): while the number of synteny breaks in the optimal order estimated for the TGD was larger in proportion to the number of loci than was the case for our previous work, POInT’s estimates of the model parameters are relatively insensitive to the exact order used (Supplemental Table 1). Similarly, the use of stringent homology criteria linked to requirements for synteny yield a set of DCS blocks that are very well supported across all eight genomes and represent a conservative set of loci with which to study the resolution of the TGD (*Methods*).

**Figure 1:**
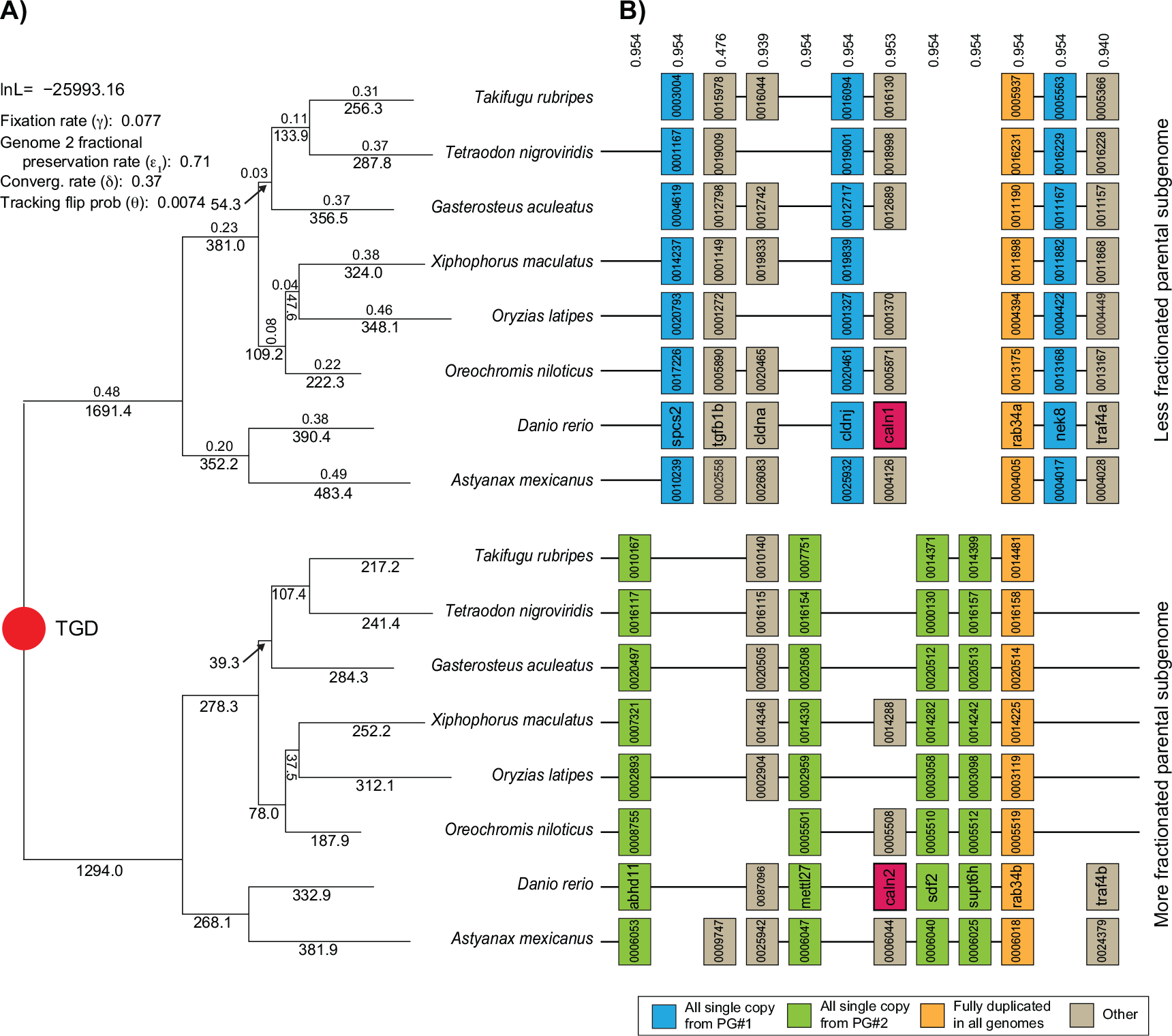
Resolution of the TGD through ohnolog losses. **A)** Shown is the assumed phylogeny of the eight species analyzed (see *Methods*). The TGD induces two mirrored gene trees, corresponding to the genes from the less fractionated parental genome (top) and the more fractionated genome (bottom). Below the branches in each tree are POInT’s predicted number of gene losses along that branch for the parental genome in question. Above the branches in the upper tree are POInT’s branch length estimates, namely the α parameter in Figure 2 and corresponding to the overall estimated level of gene loss on that branch (larger α implies a greater number of losses relative to the total number of surviving ohnologs at the start of the branch). In the upper left are POInT’s parameter estimates (γ,ε_1_,δ) for the WGD-*bc*^*nbn*^*f* model (see Figure 2). **B)** An example region of the eight genomes, showing the blocks of DCS. For all species except zebrafish, truncated Ensembl gene identifiers are given; for zebrafish gene names are shown. The numbers above each column gives POInT’s confidence in the orthology relationship shown, relative to the *2*^8^-1(=255) other possible orthology relationships. Genes are color-coded based on the pattern of ohnolog survival in the eight genomes. A pair of ohnologs expressed in the zebrafish retina are shown in magenta.

### Ohnolog fixation, biased fractionation and convergent losses are all observed after the TGD

POInT uses the copy-number status of each DCS locus in each genome, which is either duplicated (states U, F or C_1_/C_2_ in Figure 2) or single-copy (states S_1_ and S_2_), to model the resolution of a WGD along a phylogeny in a manner analogous to models of DNA sequence evolution. This computation simultaneously considers phylogenetic topology, orthology relations and copy-number states, and, unlike all other model-based approaches to gene family evolution, uses synteny data, meaning that the orthology inferences for linked loci are mutually reinforcing. Note that POInT does not use the DNA sequences of the genes in question in its computations once the homology estimation step is complete. The model parameters are instead estimated via maximum likelihood from the observed set of single-copy and duplicated loci and their synteny relationships as previously described (Conant and Wolfe 2008). As part of this computation, POInT phases the synteny blocks into to groups of orthologs (top and bottom blocks in Figure 1) by summing over all 2^n^ possible orthology states at each DCS locus (where *n* is the number of genomes), resulting in inferences such as those shown in Figure 1. Note that this orthology inference procedure accounts for the reciprocal gene losses that can create single-copy paralogs in taxa sharing a WGD (Scannell, et al. 2006); it is entirely distinct from more common orthology inference approaches that do not account for shared polyploidies (Li, et al. 2003; Jensen, et al. 2008).

**Figure 2:**
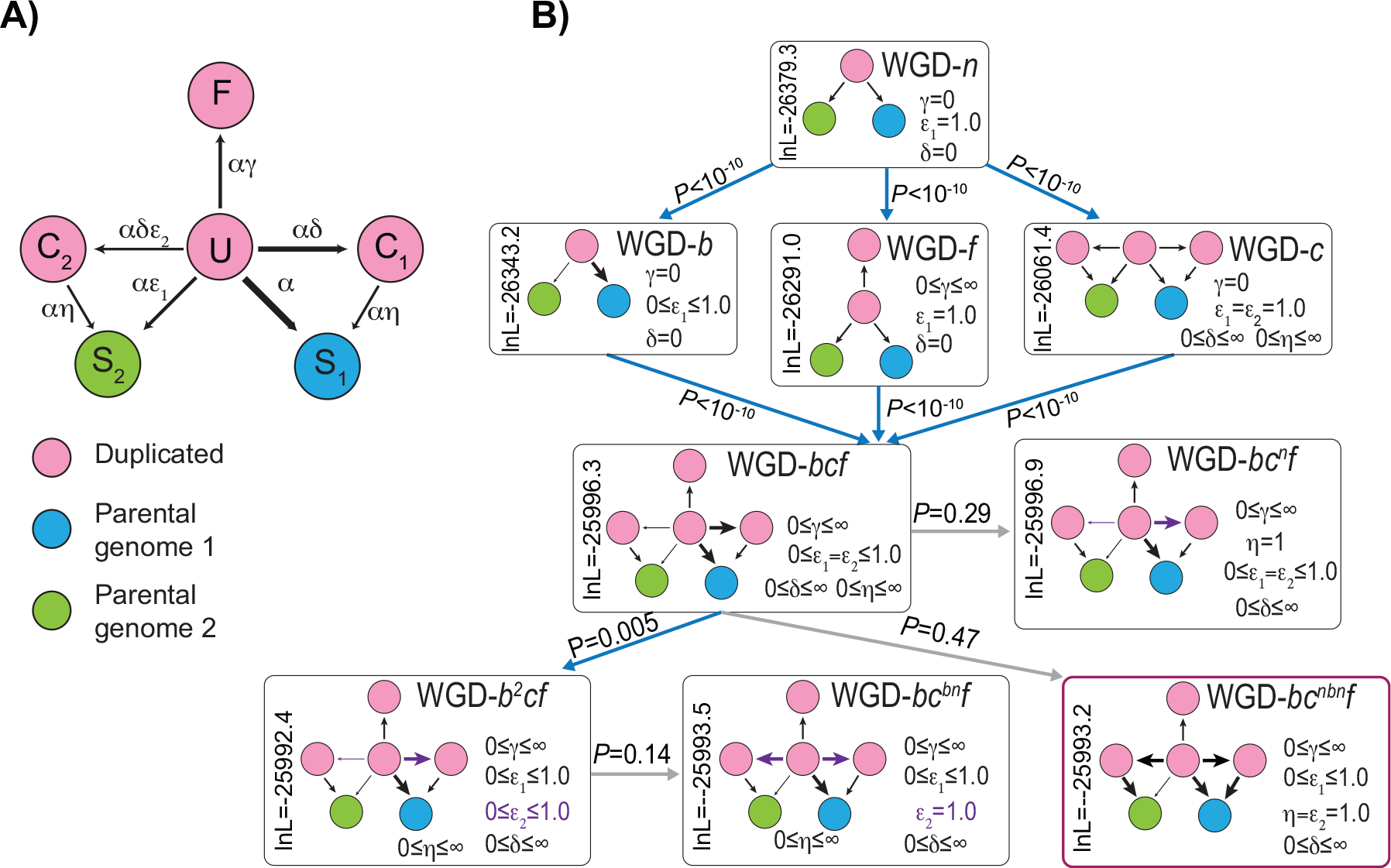
Testing nested models of post-WGD ohnolog evolution. **A)** Model states and parameter definitions for the set of models considered. **U**, **C**_**1**_, **C**_**2**_ and **F** are duplicated states, while **S_1_** and **S_2_** are single-copy states (see *Methods*). **C_1_** and **S_1_** are states where the gene from the less-fractionated parental genome will be or are preserved, and **C**_**2**_ and **S**_**2**_ the corresponding states for the more-fractionated parental genome. The fractionation rate ε (the probability of the loss of a gene from the less fractionated genome relative to the more fractionted one) can either be the same for conversions to **C**_**1**_ and **C**_**2**_ as it is for **S**_**1**_ and **S**_**2**_ (ε_1_=ε_2_) or it can differ (see **B**). **B)** Testing nested models of WGD resolution. The most basic model (top) has neither biased fractionation nor duplicate fixation nor convergent losses. Adding any of these three processes improves the model fit (second row; *P*<10^−10^). Adding the remaining two processes also improves the fit in all three cases (WGD-*bcf* model in the third row; *P*<10^−10^). However, there is no significant evidence that the ε_2_ parameter is significantly different from 1.0 (WGD-*bc^bn^f* verses WGD-*b*^2^*cf* or WGD-*bc^nbn^f* verses WGD-*bcf*), implying no biased fractionation in the transitions to states **C_1_** and **C**_**2**_. Likewise, there is no evidence that the η parameter is significantly different from 1.0 (WGD-*bc*^*n*^*f* verses WGD-*bcf* or WGD-*bc*^*nbn*^*f* verses WGD-*bcf*), meaning that losses from **C**_**1**_ and **C**_**2**_ occur at similar rates as do losses from **U**. Hence, the WGD-*bc*^*nbn*^*f* model is best supported by these data and is used for the remaining analyses.

I fit nested models of WGD evolution to the estimated ancestral order with the highest ln-likelihood under the WGD-*bc*^*nbn*^*f* model (Figure 2) in order to assess which of three processes observed after other WGD events were also detected after the TGD. The first process is duplicate fixation, meaning that some ohnolog pairs persist across the phylogeny longer than would be expected. The second process is biased fractionation, meaning that ohnolog losses favor one of the two parental genomes (defined as “Parental Genome 1” in Figure 2), and the third is the presence of convergent losses. These losses represent overly frequent parallel losses of the same member of the ohnolog pair on independent branches of the phylogeny. No matter what the order that these three phenomena are added to the duplicate loss model, all three are independently statistically significant (*P* <10^−10^; Figure 2). From these analyses, I obtained lists of surviving ohnolog pairs from zebrafish (*Dr_Ohno_all* and *Dr_Ohno_POInT*, for all zebrafish ohnologs and zebrafish ohnologs also found syntenically in other genomes, respectively; see *Methods)*, the corresponding single-copy gene sets (*Dr_Sing_all* or *Dr_Sing_POInT*) and a set of early and late ohnolog losses (e.g., losses along the root and zebrafish tip branches of Figure 1A: *POInT_RootLosses* and *POInT_DrLosses*, respectively, see *Methods*).

The results in Figure 2 notwithstanding, because each synteny block will have some variation in loss patterns, it is possible that the estimates of the strength of biased fractionation were artifacts of stochastic variation in the loss patterns in those different blocks. In our previous work (Emery, et al. 2018), we visually assessed and rejected this possibility, but there was a degree of subjectivity to that approach. Here, I instead used POInT to simulate sets of 8 genomes under a model without biased fractionation (WGD-*f*). For each simulation, I then estimated the value of the ε parameter under a model with such a bias (WGD-*bf*, see *Methods*), allowing me to assess what degree of spurious bias might be induced by our approach. The level of biased fractionation seen after the TGD is inconsistent with purely stochastic variation (*P<*0.01, Figure 3), strongly supporting the conclusion that biased fractionation occurred after the TGD.

**Figure 3:**
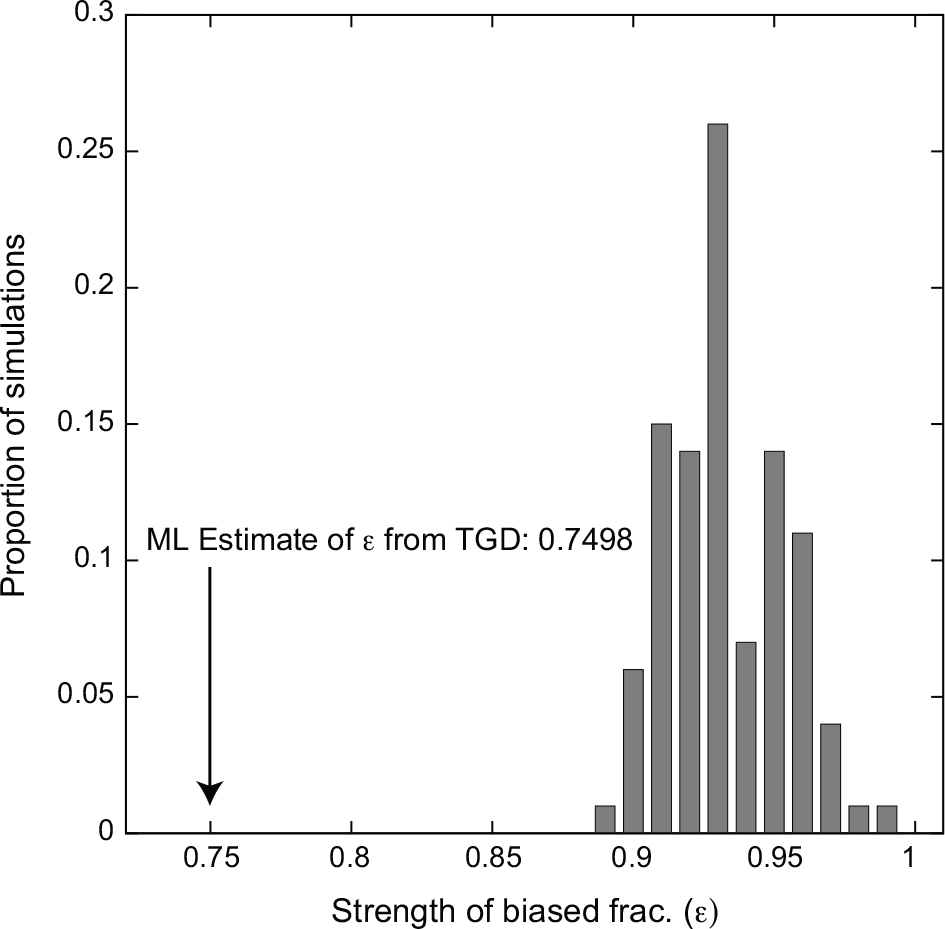
The estimated value of the biased fractionation parameter ε in the real teleost genomes (WGD*-bf* model, arrow, see *Methods*) is significantly different than those estimated from simulated genomes where biased fractionation was explicitly not included in the model (e.g., simulated ε=1.0, bars). Estimates of ε from these 100 simulations are always less than 1.0 because the model fits stochastic variations in the preservation patterns as potential biased fractionation. However, this stochastic variation never yields estimates of ε as small as seen in the real dataset (*P*<0.01).

### Ohnolog pairs are unusually rare amongst genes expressed in the earliest stages of development

As mentioned, the TGD affords the opportunity to study the effects of WGD on the evolution of developmental pathways. We had speculated that genes used in the earliest stages of development might be overly likely to be preserved in duplicate after WGD because the noise buffering effects of gene duplication might be beneficial at such times (Raser and O’Shea 2005; Pires and Conant 2016). However, such is not the case: genes with mRNAs present in the zygote were much *less* likely to be preserved as ohnolog pairs than genes first expressed later in development (Figure 4). I wondered if this observation might be driven by a dearth of ohnolog pairs among those genes where mRNA transcribed from the material genome is used in the early embryo (maternal mRNAs), since such sex-biased expression patterns might favor early gene losses. Aanes et al., (2011) have partitioned the mRNAs present in the earliest stages of zebrafish development into three groups: maternal transcripts and those seen prior to and after the mid-blastula transition. As Table 1 shows, there is some deficit of ohnologs amongst the maternally-expressed genes, but the significance of this deficit depends on the ohnolog set used, and there is no excess of early duplicate losses among this set. In contrast, the genes expressed from the embryonic genome prior to the MBT are strongly depleted in ohnologs and the single-copy genes in question are more likely to have returned to single copy along the root branch than expected (Table 1). There is then relatively little signal of ohnolog excess or deficit amongst the genes expressed later in development (post-MTB).

**Figure 4:**
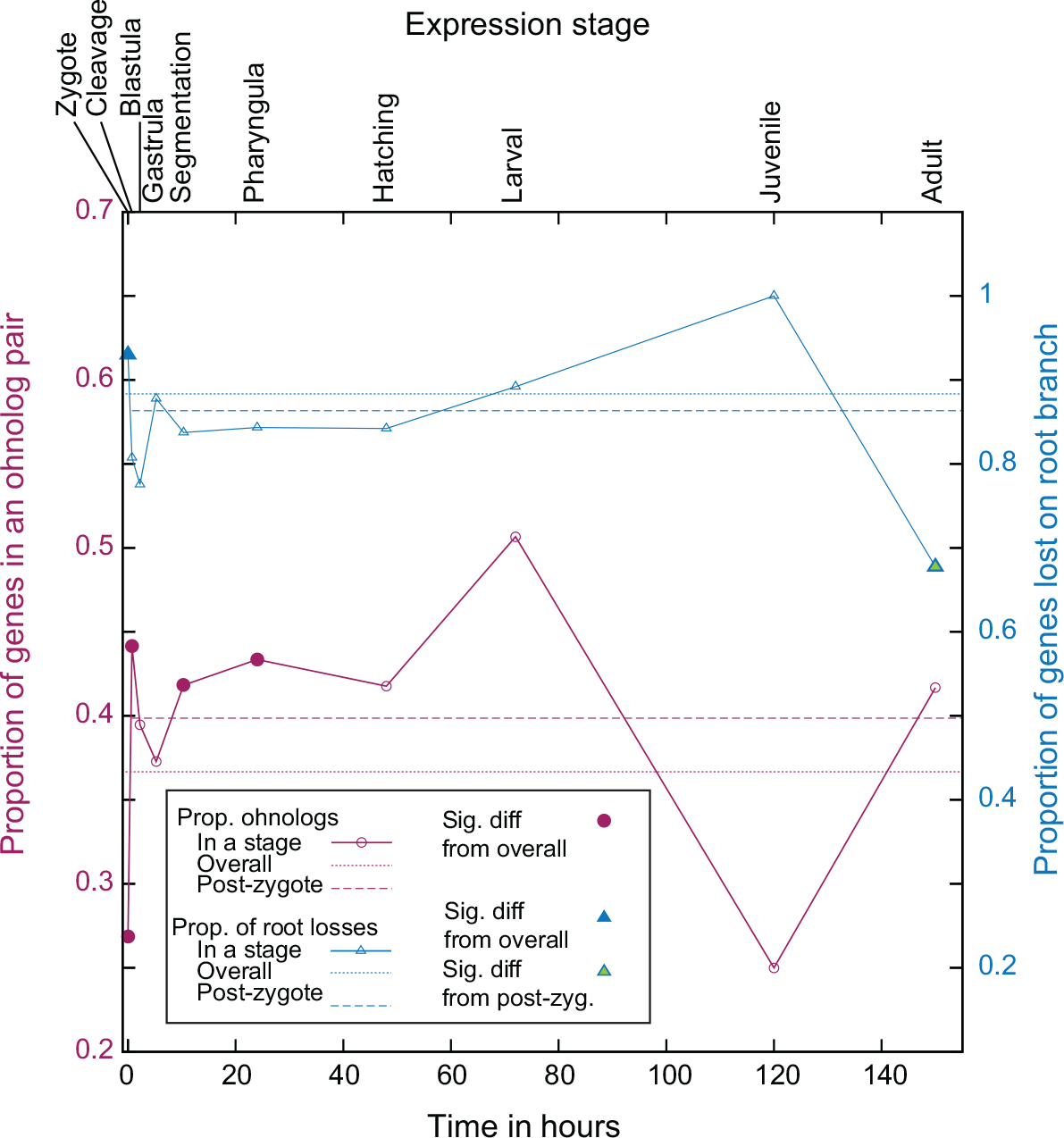
Timing of gene expression in development and patterns of ohnolog loss and retention. On the *x*-axis is a timeline of zebrafish development from ZFIN (Howe, Bradford, et al. 2013), with the relevant stage names indicated at the top. The trendline in red indicates the proportion of zebrafish genes with an ohnolog partner expressed at that stage (relative to total number of zebrafish genes analyzed with POInT expressed at that stage). The dotted red line is the overall proportion of genes with an ohnolog partner in the POInT dataset, while the dashed line is this proportion excluding any genes expressed in the zygote (see *Methods*). Open points show no statistically distinguishable difference from the overall proportion (chi-square test with an FDR correction, P>0.05; Benjamini and Hochberg 1995). Red-filled points are significantly different from this overall mean (*P*≤0.05). Trendlines in blue show similar values comparing the set of genes that POInT predicts were returned to single copy along the root branch of Figure 1 (confidence ≥ 0.85) to those only returned to single-copy along the tip branch leading to zebrafish. Hence, the right *y*-axis gives the proportion of losses that occurred along the root branch (relative to the sum of that number and the number of losses along the zebrafish branch). The dotted blue line is the overall proportion of genes returned to single-copy on the root branch (scaled as just described) while the dashed line is this proportion excluding any genes expressed in the zygote (see *Methods*). Open points are not statistically different from the overall proportion (chi-square test with an FDR correction, P>0.05; Benjamini and Hochberg 1995). Blue-filled points are significantly different from this mean (*P*≤0.05), while green filled points are also different from the mean seen when zygotic-expressed genes are excluded (*P≤*0.05).

**Table 1.** Expression timing and fate of TGD-produced ohnologs

| Expression cluster<sup>a</sup> | 1<sup>st</sup> gene set | 2<sup>nd</sup> gene set | Prop. of 1<sup>st</sup> set in cluster<sup>b</sup> | Prop. of 2<sup>nd</sup> set in cluster<sup>b</sup> | P<sup>c</sup> |
| --- | --- | --- | --- | --- | --- |
| Maternal transcripts<sup>d</sup> | *Dr_Ohno_all*<sup>e</sup> | *Dr_Sing_all*<sup>e</sup> | 0.03<br>(116/4279) | 0.04<br>(484/11616) | **2.4x10<sup>-5</sup>** |
|  | *Dr_Ohno_POInT*<sup>f</sup> | *Dr_Sing_POInT*<sup>f</sup> | 0.03<br>(81/2552) | 0.04 (193/4408) | **0.015** |
|  | *POInT_RootLosses*<sup>g</sup> | *POInT_DrLosses*<sup>g</sup> | 0.05<br>(103/1894) | 0.03 (8/250) | 0.18 |
| Pre-MBT transcripts<sup>h</sup> | *Dr_Ohno_all* | *Dr_Sing_all* | 0.10<br>(435/4279) | 0.15<br>(1709/11616) | **1.2x10<sup>-13</sup>** |
|  | *Dr_Ohno_POInT* | *Dr_Sing_POInT* | 0.11<br>(284/2552) | 0.17 (762/4408) | **3.9x10<sup>-12</sup>** |
|  | *POInT_RootLosses* | *POInT_DrLosses* | 0.19<br>(351/1894) | 0.12 (31/250) | **0.022** |
| Zygotic transcripts<sup>i</sup> | *Dr_Ohno_all* | *Dr_Sing_all* | 0.06<br>(250/4279) | 0.05<br>(573/11616) | **0.024** |
|  | *Dr_Ohno_POInT* | *Dr_Sing_POInT* | 0.06<br>(142/2552) | 0.05 (216/4408) | 0.25 |
|  | *POInT_RootLosses* | *POInT_DrLosses* | 0.05 (92/1894) | 0.05 (12/250) | >0.95 |

**a:** Three expression clusters drawn from Aanes et al., (Aanes, et al. 2011).
**b:** Proportion of all genes in the 1<sup>st</sup> set (see left) that were observed to be expressed in the cluster in question, with the total number of expressed genes over the total number of genes in that set given in parentheses.
**c:** P-value for the hypothesis test of equal proportion of genes in both sets falling into the expression cluster (chi-square test with 1 degree of freedom)
**d:** Genes determined by Aanes et al., to have been expressed in the developing embryo from maternally-derived transcripts.
**e:** Comparison of all identified zebrafish ohnologs to all zebrafish single-copy (with respect to the TGD) genes. See *Methods* for further details.
**f:** Comparison of all zebrafish ohnolog pairs found in the 8-species POInT analysis to the corresponding zebrafish single-copy (with respect to the TGD) genes. See *Methods* for further details.
**h:** Genes determined by Aanes et al., to have been expressed in the developing embryo prior to the mid-blastula transition (<3.5 hours post-fertilization).
**i:** Genes determined by Aanes et al., to have been expressed in the developing embryo only after the mid-blastula transition (>3.5 hours post-fertilization).

### GO analyses show similar patterns of ohnolog loss and retention as seen in other ancient polyploids

I used the PANTHER classification system (Mi, et al. 2017) to look for over and under-represented functions among the surviving ohnologs (and among the early ohnolog losses) in zebrafish. Supplemental Table 3 gives the complete list of significantly over and under-represented GO terms across the three hierarchies (molecular function, biological process, and cellular compartment). Here I discuss some notable results from the *Dr_Ohno_POInT* to *Dr_Sing_POInT* comparison.

Some of the Molecular Function terms over-represented among surviving ohnologs mirror results from other polyploids, such as “translation regulator activity” (*P=*0.01), “kinase activity” (*P=*0.008) and “sequence-specific DNA binding transcription factor activity,” (*P=*0.02). Many of the Biological Process terms found to be over-represented involve various aspects of nervous system function: “nervous system development” (*P<*10^−5^), “neuron-neuron synaptic transmission” (*P<*10^−5^), “synaptic vesicle exocytosis” (*P=*0.0014) and “sensory perception” (*P<*10^−4^).

I was particularly interested to see if the *under*-represented terms might shed any light on the relative absence of ohnologs among the mRNAs present in the earliest stages of development. And indeed, the four most statistically under-represented Biological Process (excepting “Unclassified”) terms among the surviving ohnologs were for “DNA metabolic process,” “translation,” “tRNA metabolic process” and “RNA metabolic process” (*P<*10^−4^ for all), while the four most significantly under-represented Molecular function terms (again excepting “Unclassified”) were “methyltransferase activity,” “structural constituent of ribosome,” “nuclease activity” and “nucleotidyltransferase activity” *(P<*10^−3^ for all). Since the earliest cell divisions in the embryo do not involve cell-type differentiation, the over-abundance of single-copy genes with roles in basic cellular processes (which would be needed even prior to such differentiation) are in accord with the expression timing results above.

### Ohnolog pairs are unusually abundant in certain nervous and sensory tissues

Using ZFIN data (Howe, Bradford, et al. 2013) on the anatomical locations of gene expression, I asked whether any embryological tissues had more or fewer members of ohnolog pairs expressed in them than expected, given the number of single-copy genes active in these same locations. Relative to the corresponding single copy genes (*Dr_Sing_POInT*), ohnologs (*Dr_Ohno_POInT*) are excessively likely to be expressed in the brain, diencephalon and epiphysis of the segmentation stage, (10.33-24 hours) and in the olfactory epithelium, retinal ganglion cell layer, and the retinal inner nuclear layer of the pharyngula stage (24-48 hours, P<0.05, chi-square test with FDR multiple test correction; Benjamini and Hochberg 1995). All of these locations except the olfactory epithelium also showed a significant excess of expressed ohnologs relative to single copy genes when the full set of zebrafish ohnologs were used (*Dr_Ohno_all* verses *Dr_Sing_all*, Supplemental Table 2). One concern with this analysis might be that the data in ZFIN are biased toward surviving ohnologs due to the genes researchers have sought to localize in the embryo: however this does not appear to be the case: 58% of ohnologs (*Dr_Ohno_POInT*) were identified in at least one anatomical location, which is actually less than the 61% of the single-copy genes (*Dr_Sing_POInT*) so identified.

### The TGD and the organization of the teleost retina

The overrepresentation of ohnologs in genes expressed in parts of the retina was intriguing because there are suggestions in the literature that the complex mosaic organization of teleost retinae (Lyall 1957; Engström 1960; Stenkamp and Cameron 2002) might be an innovation due to the TGD (Braasch and Postlethwait 2012; Sukeena, et al. 2016). I conducted a GO analysis of all ohnologs and single-copy genes expressed in either the ganglion or inner cell layers of the retina at the pharyngula stage of development. No terms associated with biological process were over-represented in either tissue, and no terms associated with molecular function were over-represented in the inner cell layer. However, for the ganglion layer, the term “transmembrane transporter activity” was significantly overabundant among the surviving ohnologs (*P*=0.044 after FDR correction). Interestingly, the expression of duplicated genes from the TGD in these locations are probably not specific to zebrafish: the *only* two GO biological process terms that are *under* represented among the genes returned to single-copy along the root branch of Figure 1 (e.g., genes that survived in duplicate at least to the first post-TGD speciation) are “synaptic transmission” and “cell-cell signaling,” while the single Cellular Compartment term similarly under represented is “neuron projection” (Supplemental Table 3). Collectively, these results suggest that the duplicated genes created by the TGD were likely involved in subsequent evolution changes in neuronal development, accounting for their retention as ohnologs across the teleost phylogeny.

### Surviving TGD ohnologs are less likely to be essential

I compared the proportion of phenotyped genes with surviving ohnologs judged to be essential in zebrafish to the same proportion among those genes without surviving ohnologs: the genes with ohnologs show a reduced propensity to be essential (Table 2, see *Methods* for details). Importantly, this effect does not appear to be a result of any intrinsic features of these genes: when examining the two groups in the unduplicated outgroup mouse, I find that that single-copy mouse orthologs of the duplicated and the unduplicated zebrafish genes have similar essentiality in that animal. However, I also note that this effect is not a strong one: when I examined the smaller set of ohnologs with support across the eight genomes (*Dr_Ohno_POInT* verses *Dr_Sing_POInT*), the proportions shown in Table 2 are nearly identical, but the effect is non-significant due to the smaller sample size (*P=*0.14, chi-square test).

**Table 2.** Essentiality and the TGD

| Essentiality data<sup>a</sup> | Prop. of phenotyped genes with an ohnolog that are essential<sup>b</sup> | Prop. of phenotyped genes without an ohnolog that are essential<sup>c</sup> | *P*<sup>d</sup> |
| --- | --- | --- | --- |
| Zebrafish<sup>e</sup> | 0.062 (6/97)<sup>f</sup> | 0.145 (46/318)<sup>f</sup> | **0.048** |
| Mouse<sup>g</sup> | 0.556 (42/72)<sup>h</sup> | 0.506 (161/318)<sup>h</sup> | 0.53 |

**a:** We compared the essentiality of genes with and without surviving TGD-produced ohnologs in zebrafish for their essentiality in zebrafish and the essentiality of their single-copy orthologs from mouse.
**b:** Proportion of genes with a ZFIN phenotype with a surviving TGD-produced ohnolog that are essential in the dataset in question (set *Dr\_Ohno\_all*).
**c:** Proportion of genes with a ZFIN phenotype without a surviving TGD-produced ohnolog that are essential in the dataset in question (set *Dr\_Sing\_all*).
**d:** *P*-value for the hypothesis test of equal proportion of essential genes in *Dr\_Ohno\_all* vs *Dr\_Sing\_all*.
**e:** Essentiality defined as genes in the ZFIN database (Howe, Bradford, et al. 2013) phenotyped as “lethal,” “dead” or “inviable.”
**f:** Numbers in the parenthesis give the number of essential genes over the total number in the set.
**g:** Essentiality defined by the International Mouse Phenotyping Consortium’s list of essential mouse genes (Koscielny, et al. 2013; Dickinson, et al. 2016).
**h:** Numbers in the parenthesis give the number of essential genes over the total number in the set. Note that ohnolog pairs in zebrafish are by definition single-copy in gar and mouse, accounting for the smaller number of comparisons in the ohnolog column for the mouse analysis.

### TGD ohnologs lie in connected parts of the zebrafish metabolic network

I examined the position of the ohnolog pairs in the published zebrafish metabolic network (Bekaert 2012). Ohnologs are more likely to be members of this network than are single copy genes (*P=*0.0005 and *P=*0.025 for *Dr_Ohno_all* verses *Dr_Sing_all* and *Dr_Ohno_POInT* verse *Dr_Sing_POInT*, respectively). Ohnolog pairs also occupy more connected parts of this network (e.g., they have higher mean degree; Table 3), though they do not different from the single-copy genes with respect to other network statistics.

**Table 3.** The TGD and the zebrafish metabolic network

| Network statistic<sup>a</sup> | Ohnolog datasets compared<sup>b</sup> | Mean ohnolog value<sup>c</sup> | Mean single-copy gene value<sup>d</sup> | *P*<sup>e</sup> |
| --- | --- | --- | --- | --- |
| Node degree<sup>f</sup> | *Dr_Ohno_all/Dr_Sing_all*<sup>g</sup> | 30.9 | 21.4 | **0.002** |
|  | *Dr_Ohno_POInT Dr_Sing_POInT*<sup>h</sup> | 31.5 | 23.2 | **0.032** |
| Avg. clustering coeff.<sup>i</sup> | *Dr_Ohno_all/Dr_Sing_all*<sup>g</sup> | 0.78 | 0.77 | >0.5 |
|  | *Dr_Ohno_POInT Dr_Sing_POInT*<sup>h</sup> | 0.77 | 0.76 | >0.5 |
| Mean # shortest paths<sup>j</sup> | *Dr_Ohno_all/Dr_Sing_all*<sup>g</sup> | 18020 | 12904 | 0.07 |
|  | *Dr_Ohno_POInT Dr_Sing_POInT*<sup>h</sup> | 19280 | 14132 | 0.18 |

**a:** Network statistic for which the ohnologs and single copy genes were compared.
**b:** Derivation of the set of ohnologs and single-copy genes.
**c:** Mean value of the statistic in question for the ohnolog pairs (ohnolog pairs were merged and averaged prior to computing the global average).
**d:** Mean value of the statistic in question for the single-copy genes.
**e:** *P*-value for the hypothesis test of equal mean statistic value for the ohnologs and single-copy genes (Network randomization test; *Methods*).
**f:** Number of edges per network node.
**g:** Comparison of all identified zebrafish ohnologs to all zebrafish single-copy (with respect to the TGD) genes. See *Methods* for further details.
**h:** Comparison of all zebrafish ohnolog pairs used in the 8 species POInT analysis to the corresponding zebrafish single-copy (with respect to the TGD) genes. See *Methods* for further details.
**i:** Ratio of the number of edges between each triplet of nodes to the maximum number of such connections possible (Watts and Strogatz 1998).

## Discussion

Polyploidies of widely differing ages are ubiquitous across the tree of life (Van de Peer, et al. 2017), yet many of the detailed studies of its genome-wide effects have focused on recent events. Thus, while we know quite a bit about the fate of individual ohnolog pairs surviving from events such as the TGD and the vertebrate 2R events (Force, et al. 1999; McLysaght, et al. 2002; Steinke, et al. 2006; Makino and McLysaght 2010; Guo, et al. 2012; Kollitz, et al. 2014; Moriyama, et al. 2016; Xie, et al. 2016), we do not currently know as much about whether the patterns of genome evolution, such as adherence to the DBH and the occurrence of biased fractionation, seen after recent polyploidies, also apply to these more ancient events. Existing data should also be interpreted with caution as the methods used to identify the relics of ancient WGDs are subject to bias. Hence, Inoue et al.,’s estimates of the timing of ohnolog losses after the TGD differ from those presented here (Inoue, et al. 2015), with their estimates of the proportion of losses along the root branch being >1.5 greater than that estimated with POInT, with an average of only 21% as many proportional losses inferred along the tip branches as POInT predicts. The reason for the discrepancy is likely that Inoue et al.,’s method does not phase post-WGD orthologs. Without such phasing, independent losses in different lineages will be mistaken for shared losses, leading to the type of over-estimates of initial losses that was apparently observed.

The data shown here support a role for the DBH in resolving the TGD: the location of ohnologs in the zebrafish metabolic network is similar to the pattern seen in the network of the polyploid plant *Arabidopsis thaliana* (Bekaert, et al. 2011) and the classes of ohnologs retained follow the predictions of the DBH (Freeling 2009; Conant, et al. 2014). However, further work will be needed to assess whether these surviving ohnologs with high interaction degree are still be maintained by selection on relative dosage or if some other force is now at work (Conant, et al. 2014).

Likewise, the TGD appears to have been an allopolyploidy, because there is strong evidence for biased fractionation and no known mechanism by which autopolyploidy could generate such biases. This result is also consistent with studies in yeast and plants (Garsmeur, et al. 2013; Marcet-Houben and Gabaldon 2015; Emery, et al. 2018). Perhaps more unexpectedly, the pattern of early gene losses after the TGD, namely among genes for the core of the cellular machinery, recall patterns seen in plants and yeast, where processes such as DNA repair were rapidly returned to single copy after polyploidy (De Smet, et al. 2013; Conant 2014). Indeed, “DNA repair” is a highly under-represented term (*P<*10^−3^) among the zebrafish TGD ohnologs, though not one of the top 4 listed above. De Smet et al., have argued that these loss patterns suggest selection to return genes with these types of function to single copy. What was less clear from previous work is that the dearth of ohnologs among these basic processes would correspond to an excess of them involved in other processes such as multicellular development. Hence, polyploidy in multicellular organisms might concentrate its effects in such developmental processes (Holland, et al. 1994).

In this vein, the over-abundance of ohnologs expressed in the developing retina is very interesting because recent work in the spotted gar strongly suggests that the mosaic organization of teleost retainae (Lyall 1957; Engström 1960; Stenkamp and Cameron 2002) represents a morphological innovation whose evolutionary appearance was coincident with the TGD (Sukeena, et al. 2016). Not only are ohnologs over-represented in genes expressed in some of the retinal layers, but a GO analysis suggested that many of these duplicated genes function as transmembrane transporters. Several analyses have suggested that cell-to-cell communication in the early stages of retinal development may drive the mosaic organization (Stenkamp and Cameron 2002; Raymond and Barthel 2004), and such transmembrane proteins are obvious candidates for such communication. Moreover, there may be other neurological innovations due to the TGD, if the excess of ohnologs in nervous tissues and with nervous system functions is a guide.

While duplicate genes can provide a “backup” for each other in response to gene knockout, this effect is expected to degrade as the duplicate pair ages (Gu, et al. 2003), making the apparent rarity of essential genes among the ohnolog pairs a bit surprising, given the TGD’s age. However, gene essentiality and gene duplications interact with each other in a complex way. On the one hand, a gene’s propensity to duplicate is associated with whether or not it is essential: small scale duplications favor less essential genes (Woods, et al. 2013), but post-WGD evolution appears to neither favor nor disfavor the retention of (formerly) essential genes after WGD (Wapinski, et al. 2007; Deluna, et al. 2008). Gene duplication then apparently imparts the (partial) redundancy seen in studies of yeast, nematodes and mice (Gu, et al. 2003; Conant and Wagner 2004; Makino, et al. 2009). I suspect that the combined observation of reduced essentiality of zebrafish ohnologs with no reduction in the essentiality of their single-copy mouse orthologs mostly likely represents surviving shared functions between ohnolog pairs that were preserved in duplicate due to other selective pressures.

The most general message apparent from these analyses is that polyploidy continues to shape the evolutionary trajectories of its possessors over very long time scales, both through first-order effects such as genetic robustness, and, more importantly, through the appearance of duplication-driven evolutionary innovations. Examples such as the changes in retinal structure just discussed are particularly important because they appear to be a class of innovations requiring changes in many genes at once, meaning that they may have only been feasible with the large number of duplicates induced by polyploidy. Though relatively few examples of such innovations are currently known (Conant and Wolfe 2007; Merico, et al. 2007; van Hoek and Hogeweg 2009; Edger, et al. 2015), as our knowledge of both polyploidy and the systems biology of the cell increases, it is very likely more will be found.

## Methods

### Identifying the relics of the TGD from double-conserved synteny blocks

We have developed a pipeline for inferring shared blocks of double-conserved synteny (DCS, Figure 1) from a group of genomes sharing a WGD and a reference genome from an unduplicated relative (Emery, et al. 2018). I applied this tool to eight polyploid fish genomes, taken from the Ensembl database (Aken, et al. 2017): *Astyanax mexicanus* (Cave fish; McGaugh, et al. 2014), *Danio rerio* (Zebrafish; Howe, Clark, et al. 2013), *Takifugu rubripes* (Fugu; Aparicio, et al. 2002), *Oryzias latipes* (Medaka; Kasahara, et al. 2007) *Xiphophorus maculates* (Platyfish; Schartl, et al. 2013), *Gasterosteus aculeatus* (Stickleback; Jones, et al. 2012), *Tetraodon nigroviridis* (Jaillon, et al. 2004) and *Oreochromis niloticus* (Tilapia; Brawand, et al. 2014). The genome of *Lepisosteus oculatus* (spotted gar; Braasch, et al. 2016) was used as the unduplicated outgroup.

The pipeline has three steps. First, I performed a homology search of each polyploid genome against the gar genome with GenomeHistory (Conant and Wagner 2002). I defined a gene from a polyploid genome to be a homolog of a gar gene if it had a BLAST E-value (Altschul, et al. 1997) of 10^−8^ or smaller and was 60% or more identical to that gene at the amino acid level. I further required that the length of the genes’ pairwise alignment be 65% or more of their mean length and that the pair have nonsynonymous divergence (K_a_) less than 0.6. These parameters result in good coverage of the genomes involved: between 70% and 80% of gar genes have a homolog in each of the TGD-possessing genomes, and between 70% and 82% of genes in those genomes have a gar homolog. Nonetheless, the parameters do not overly merge gene families, with 58% to 60% of the gar genes showing only a single homolog in the TGD-possessing genomes.

This set of homologs was then the input to the second step of the pipeline: the inference of DCS blocks in each polyploid genome. This step determines which of the potentially many homologs of a given gene in gar are the ohnologs from the TGD by maximizing the number of homologs placed in the DCS blocks. We have referred to each such gar gene and its corresponding two potential products of the TGD as a “pillar” (Byrne and Wolfe 2005; Emery, et al. 2018): the resulting set of these *n* pillars is denoted *A*_1_..*A*_*n*_. Each pillar has associated with it a set of homologous genes from the polyploid genome *h*_1_…*h*_*h*_. At most two of these homologs can be assigned to the pillar’s ohnolog positions, denoted *A*_*i*_(*p*_1_) and *A*_*i*_(*p*_2_). We define *A*_*O*(*i*)_ to be the *i*^*th*^ pillar in the reordered version of this dataset. It is necessary to estimate the *A*_*O*(*i*)_s because the teleost genomes have undergone rearrangements since the TGD (Nakatani and McLysaght 2017). Using simulated annealing (Kirkpatrick, et al. 1983; Conant and Wolfe 2006), I sought the combination of homolog assignments and pillar order that maximizes the number of pillars where the genes in neighboring pillars are also neighbors in their genome (Emery, et al. 2018). Precisely, I maximized the score *s* of such a combination of homolog assignments and pillar orders:

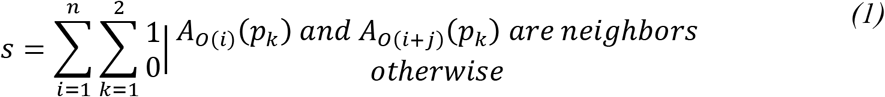

Once those inferences were complete for each of the eight polyploid genomes, I merged them, mapping between polyploid genomes using the gar genes as references. Taking an extremely conservative approach, I retained pillars only if each assigned homolog from every genome had synteny support in at least one direction. The result was 5589 pillars with at least one syntenic gene from each polyploid genome. I then again used simulated annealing to infer the optimal pillar order over all eight genomes. Because of the high degree of rearrangement, I made inferences of the optimal ordering under three different criteria. First, as previously (Emery, et al. 2018), I started with the order of the gar reference genes and sought orderings with the fewest total synteny breaks (*Naïve_Opt*). Second, I used an initial greedy search to place pillars with many neighboring genes in the eight extant teleost genomes near to each other, which reduced the number of initial breakpoints by about 30%. I then again sought an order with minimal breaks (*Greedy_Opt*). Finally, I sought an ordering that maximized the number of neighboring pillars having no synteny breaks between them in any genome and, after using this optimization criterion for several iterations, again applied the standard search for the fewest total breaks (*Global_Break_Opt*). I then used the inferred order that gave the highest likelihood of observing that WGD data under the WGD-*bc*^*nbn*^*f* gene loss model (Supplemental Table 2; see *Modeling the evolution of the TGD* below) for all further analyses.

### Quality of inferred double-conserved synteny blocks

Given the ancient nature of the TGD, it is reasonable to ask if this DCS inference protocol is sufficient for these genomes. However, despite the divergence, the mapping between the genomes possessing the TGD and spotted gar is less difficult than might be expected, with 69-71% of the genes in the teleost genomes in our final dataset having only a single gar homolog (and where that gar gene matches at most 2 genes in the genome with the TGD; Supplemental Table 1). Although I required every analyzed gene to be in synteny in Step 2 of the pipeline, the estimate of a global ancestral order requires breaking some of these synteny blocks. But this problem is not serious: >94% of the genes across all the genomes with the TGD that I analyzed are in synteny blocks in the estimated ancestral order used, with the large majority in blocks of 5 or more genes (Supplemental Table 1). I provide the synteny relationships under the inferred order for the eight genomes as supplemental data.

For comparative purposes, I also explored whether a gene tree/species tree reconciliation approach might offer improved accuracy relative to synteny-based methods. Of the 132 loci in the dataset where all eight species share preserved ohnolog pairs, there are nine loci where all of the 16 genes that are members of these ohnolog pairs show clear syntenic associations in both directions. Such positions represent the best-case scenario for gene tree-based methods, as there is no gene loss to confound the inference process. I extracted the (9×8×2=144) genes in question and made codon-preserving alignments with T-Coffee (Notredame, et al. 2000) for each of the nine loci. Using phyml (Guindon and Gascuel 2003), I inferred maximum likelihood trees from these alignments under the GTR model with 4 categories of substitution rates that followed an estimated discrete gamma distribution. For none of the nine loci was the expected pair of mirrored species trees inferred (see Figure 1). In fact, of the 18 gene trees inferred (two per ohnolog pair), only 3 matched the assumed species tree, and no other topology was seen more frequently. Hence, while it is possible to infer products of the TGD by reconciling such gene trees with a proposed species tree using tools such as NOTUNG (Chen, et al. 2000), there is little reason to believe that such an approach would perform as well as the one employed. Moreover there are good theoretical reasons to believe that genes tree inference from these genomes should be challenging: most obviously, the relationships in question are characterized by long branches and may experience significant gene conversion post-WGD (Felsenstein 1978; Evangelisti and Conant 2010; McGrath, et al. 2014; Scienski, et al. 2015).

### Modeling the evolution of the TGD

I analyzed the DSC blocks from these genomes using POInT (Conant and Wolfe 2008; Conant 2014) under several models of post-WGD duplicate loss. These models have four to six states (Figure 2): **U** (undifferentiated duplicated genes), **F** (fixed duplicate genes), **S**_**1**_ and **S**_**2**_ (single copy states) and the converging states **C**_**1**_ and **C**_**2**_. These last two states model the potential for the independent parallel losses first seen in yeast (Scannell, et al. 2007; Conant and Wolfe 2008). I used likelihood ratio tests to identify the combination of these factors best fitting the data (Figure 2; Sokal and Rohlf 1995). POInT’s orthology inferences for all pillars are given as Supplemental Data.

### Simulating genome evolution under a model where no biased fractionation occurs

We have previously described using POInT as a simulation engine for generating sample genome duplications (Conant and Wolfe 2008). Briefly, I started from a set of completely duplicated loci and the assumed gene order previously estimated. In locations where gene losses in one genome had generated a synteny break (e.g., after *caln1* in Figure 1), I extended the left contig to include the introduced duplicates. Then, using the maximum likelihood estimates of the model parameters and branch lengths under the WGD-*f* model, I generated a new set of post-WGD duplicate losses along the phylogeny of Figure 1. Finally, I applied the “Tracking flip prob.” parameter noted in Figure 1 to model POInT’s estimated errors in orthology inference, introducing new synteny breaks in the simulated genomes whenever a uniform random number was drawn with a value less than this parameter. I analyzed 100 such simulated sets of genomes with POInT under the WGD-*bf* model (e.g., biased fractionation and fixation allowed, but the δ parameter in Figure 2 set to 0) and extracted the value of ε, which is plotted in Figure 3. No simulated dataset had a value of ε as small as seen in the real dataset (*P*<0.01).

### The TGD and the teleost phylogeny

As our estimate for the phylogenetic relationships of these 8 species, we used the phylogeny of Near et al., (2012). While exhaustively examining all possible topologies was not computationally feasible, I was able to examine 4 trees that were near topological neighbors to that of Near et al,: all gave lower likelihoods of observing the genomic data than did that of Near et al., (Supplemental Figure 1). I note that the TimeTree package estimates that the first split between the eight taxa studied here occurred between 230 and 315 million years ago (Kumar, et al. 2017), consistent with the TGD’s estimated age.

### Zebrafish ohnolog and single-copy gene sets

Based on the inferences above, I defined two sets of zebrafish ohnologs and corresponding single copy genes. *Dr_Ohno_all* is the set of all ohnolog pairs that are part of DCS blocks found in the pairwise comparison of *D. rerio* to gar; *Dr_Sing_all* is the corresponding WGD loci that have returned to single copy. *Dr_Ohno_POInT* corresponds to the set of ohnologs from zebrafish for which the WGD locus/pillar in question was also identified in the other seven polyploid teleost genomes, with *Dr_Sing_POInT* being the corresponding single copy set. I also defined a pair of gene sets consisting of genes that POInT predicts with high confidence (*P≥*0.85) to have been returned to single copy on the shared root branch of the phylogeny in Figure 1 (*POInT_RootLosses*) and a corresponding set predicted with the same confidence to have been lost only on the branch leading to the extant *D. rerio* (e.g., after the split of zebrafish and cavefish; *POInT_DrLosses*).

### Gene expression timing and WGD

I extracted the earliest developmental stage at which each zebrafish gene’s transcript has been observed and the corresponding time of expression in terms of hours post-fertilization from the ZFIN database (Howe, Bradford, et al. 2013). From the same database, I also extracted all non-adult anatomical locations at which each gene’s transcript had been detected. For each developmental stage and location, I used a chi-square test with a false-discovery rate correction (Benjamini and Hochberg 1995) to test for differences in the proportion of ohnologs and non-ohnologs (*Dr_Ohno_all* vs *Dr_Sing_all* and *Dr_Ohno_POInT vs Dr_Sing_POInT*) expressed at that location. I similarly compared the proportion of single copy genes in each location and stage that were early and late losses (*POInT_RootLosses* verses *POInT_DrLosses*). For the anatomical tests, any gene expressed in the zygote was omitted from the analysis to avoid having the strong bias against ohnologs in this stage give rise to spurious associations.

Aanes et al., (2011) have partitioned gene expression in the early zebrafish embryo into three groups: genes expressed from inherited maternal transcripts, genes expressed from the embryo’s genome prior to the midblastula transition (pre-MTB) and genes expressed first in the zygotic stage (e.g., post-MTB). Using these gene lists, I compared the frequency of ohnologs and single-copy (*Dr_Ohno_all* vs *Dr_Sing_all* and *Dr_Ohno_POInT vs Dr_Sing_POInT*) genes in each, again using a chi-square test, as well as the proportion of root losses and tip losses (*POInT_RootLosses* vs *POInT_DrLosses*) with the same approach (Supplemental Table 3).

### GO analyses

To understand how gene function may have shaped the resolution of the TGD, I used the Gene List Analysis tool from the PANTHER classification system (version 13.1; Mi, et al. 2017) to find over or under-represented Gene Ontology (GO) terms associated with the surviving ohnologs (*Dr_Ohno_all* compared to *Dr_Sing_all* and *Dr_Ohno_POInT* to *Dr_Sing_POInT*) and the early verses late ohnolog losses (*POInT_RootLosses* compared to *POInT_DrLosses*). In each case, I asked whether there were any biological processes, molecular functions, or cellular component ontology terms that were significantly over or under-represented on the first list, using Fisher’s exact test with an FDR multiple test correction (Mi, et al. 2013; Mi, et al. 2017). Lists of all significantly enriched terms for any comparison are given as Supplemental Table 4.

### Gene essentiality and the TGD

From ZFIN (Howe, Bradford, et al. 2013), I extracted all genes with known phenotypes, as well as the subset of those genes with phenotypes described as “lethal,” “dead” or “inviable:” hereafter I note this second set as the “essential genes.” I compared the proportion of phenotyped ohnologs in the essential list to the same proportion among the single copy genes. For comparative purposes, I obtained a list of essential mouse genes from the International Mouse Phenotyping Consortium (Koscielny, et al. 2013; Dickinson, et al. 2016). Using our orthology inference pipeline, I inferred the gar orthologs of these mouse genes (Conant 2009; Bekaert and Conant 2011), retrieving 10,644 gar genes with a mouse ortholog. For each gar gene with phenotype data in a mouse ortholog, we compared the proportion of genes with a surviving ohnolog in zebrafish that were essential when knocked out in mouse to the proportion of genes without a surviving zebrafish ohnolog pair that were essential (Table 2; other phenotype classes such as “subviable” were excluded).

### The TGD and the zebrafish metabolic network

I extracted an enzyme-centered metabolic network from the reconstruction of zebrafish metabolism published by Bekaert (2012). In this network nodes are biochemical reactions and edges connect pairs of nodes with a common metabolite. The 13 currency metabolites given by Bekaert (2012) were excluded from the edge computation. Each reaction was linked to one or more Ensembl gene identifiers corresponding to genes encoding enzymes catalyzing that reaction.

To test for differences in network position between the products of ohnologs and single-copy genes, I compared the two groups for their average degree (number of edges), clustering coefficient (indicating the propensity of connected nodes to have common neighbors; Watts and Strogatz 1998), and betweenness-centrality (the number of the network’s shortest paths passing through a given node; Hahn and Kern 2005). I then used randomization to assess the statistical significance of the differences. To maintain the structure introduced by the WGD, all ohnolog pairs were reduced to a single entity, which was then assigned to all nodes that products of either of the two ohnologs appeared in. These merged ohnolog products were then randomized along with the products of the single copy genes, and the differences in the three statistics for each randomized network recomputed. If less than 5% of the randomized networks had a difference as large as that observed for the real data, I concluded that there was evidence for a difference between duplicated and unduplicated genes.

## Abbreviations

WGD: whole-genome duplication
TGD: teleost genome duplication
DCS: double conserved synteny
POInT: Polyploid Orthology Inference Tool

## Acknowledgements

I would like to thank J. C. Pires, J. Thorne and X. Ji for helpful discussions and K. Dudley for computational assistance. This work was supported by the United States National Science Foundation (grant numbers NSF-IOS-1339156 and NSF-CCF-1421765).

## Supplemental Information

**Supplemental Figure 1:**
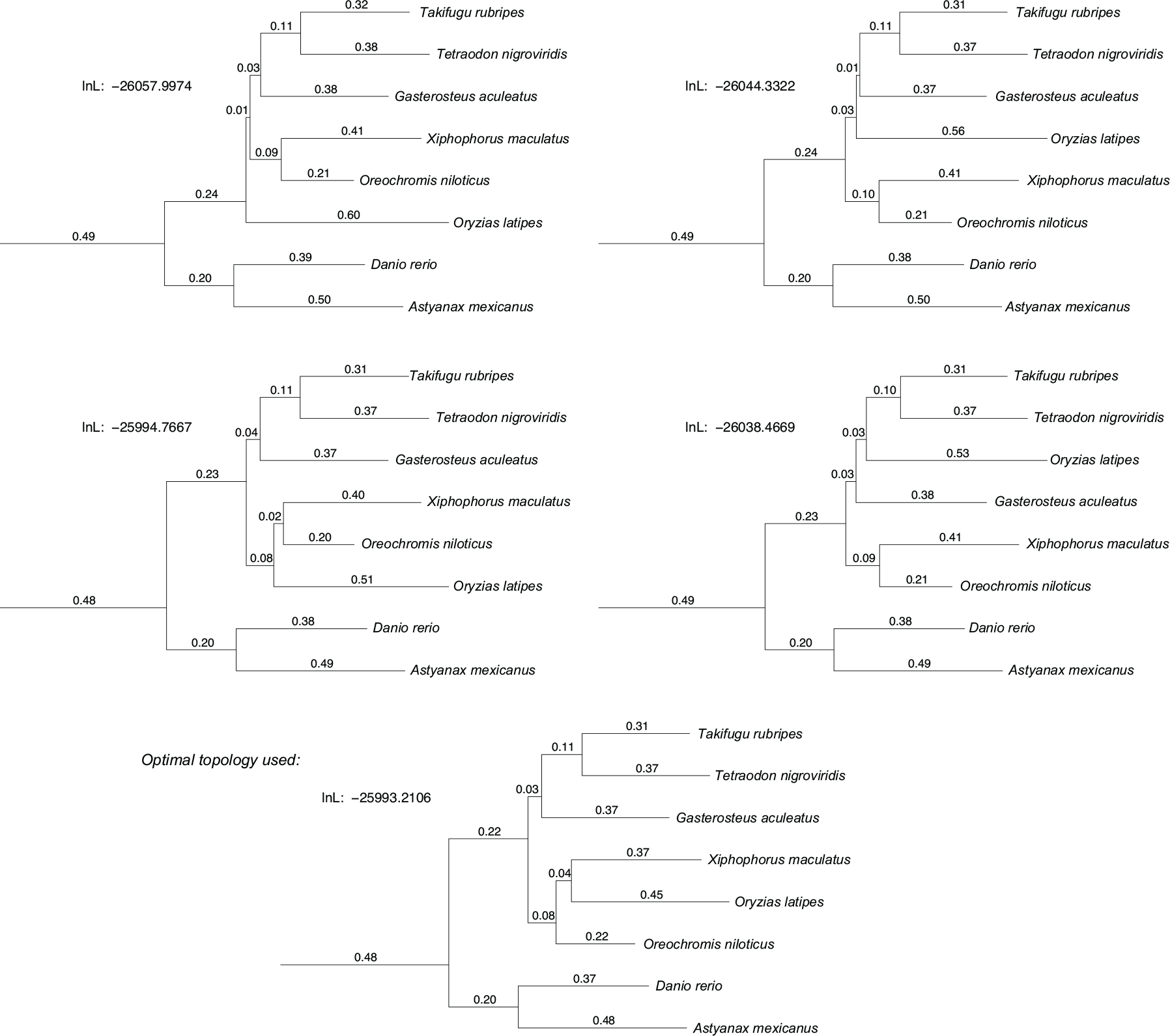
Alternative phylogenetic topologies and the TGD. I tested 4 other topologies in addition to that of Near et al., (Near, et al. 2012) using the optimal DCS block order and the WGD-*bc*^*nbn*^*f* model in POInT: shown are the induced branch lengths for these topologies as well as the corresponding model parameter estimates. For reference, the topology at the bottom is that of Near et al., used for all other analyses.

**Supplemental Table 1:** Quality of inferred DCS blocks across the eight genomes.

| Species | Total genes in dataset<sup>a</sup> | % of genes with 1:1/1:2 homology to gar<sup>b</sup> | % genes in synteny blocks<sup>c</sup> | % genes in synteny blocks w/ >5 genes<sup>d</sup> | # of genes not in synteny blocks with ambig. homology<sup>e</sup> |
| --- | --- | --- | --- | --- | --- |
| *Astyanax mexicanus* | 6707 | 71.4 | 94.0 | 80.2 | 116 |
| *Danio rerio* | 6849 | 69.3 | 95.3 | 85.1 | 97 |
| *Gasterosteus aculeatus* | 6799 | 69.6 | 99.2 | 98.0 | 22 |
| *Oreochromis niloticus* | 6936 | 69.6 | 99.4 | 97.8 | 13 |
| *Oryzias latipes* | 6601 | 69.9 | 99.3 | 97.3 | 12 |
| *Takifugu rubripes* | 6725 | 70.4 | 99.0 | 96.6 | 34 |
| *Tetraodon nigroviridis* | 6660 | 70.0 | 98.7 | 96.4 | 41 |
| *Xiphophorus maculatus* | 6685 | 70.1 | 98.0 | 93.4 | 41 |

**a:** Number of genes from the genome placed into the shared DCS blocks used for POInT analyses (see *Methods*).
**b:** Percent of cases in the genome in question where genes in the final dataset from that genome match one and only one gene in the gar genome and the gar gene in question matches at most two genes in that genome.
**e:** Number of genes in the dataset analyzed which are not in synteny blocks in the optimal order that also have some ambiguity in their homology relationships: either a teleost gene with multiple gar homologs or where the gar gene in question has >2 homologs in this genome.

**Supplemental Table 2:** Effects of the estimated ancestral pillar order on parameter estimates from POInT.

| Optimization Method<sup>a</sup> | Breaks<sup>b</sup> | lnL<sup>c</sup> | $\gamma^d$ | $\epsilon^d$ | $\delta^d$ | $\theta^d$ |
| --- | --- | --- | --- | --- | --- | --- |
| Global_Break_Opt | 14342 | -26240.049 | 0.075 | 0.676 | 0.342 | 0.0089 |
|  | 14693 | -26170.596 | 0.074 | 0.669 | 0.339 | 0.0085 |
|  | 14787 | -26188.226 | 0.075 | 0.684 | 0.343 | 0.0084 |
|  | *14965<sup>e</sup>* | *-26080.462* | *0.074* | *0.657* | *0.341* | *0.0079* |
|  | 15148 | -26178.494 | 0.076 | 0.683 | 0.352 | 0.0076 |
|  | 15966 | -26272.204 | 0.076 | 0.728 | 0.382 | 0.0076 |
|  | 16323 | -26272.011 | 0.075 | 0.673 | 0.361 | 0.0073 |
|  | 16766 | -26323.416 | 0.074 | 0.680 | 0.379 | 0.0074 |
|  | 17523 | -26265.175 | 0.076 | 0.723 | 0.407 | 0.0066 |
|  | 21656 | -27092.183 | 0.071 | 0.634 | 0.416 | 0.0052 |
| Greedy_Opt | 13852 | -26226.784 | 0.077 | 0.701 | 0.370 | 0.0097 |
|  | 13898 | -26203.132 | 0.078 | 0.711 | 0.370 | 0.0087 |
|  | 13937 | -26103.825 | 0.077 | 0.685 | 0.356 | 0.0084 |
|  | 13974 | -26169.419 | 0.077 | 0.704 | 0.361 | 0.0087 |
|  | 14086 | -26136.803 | 0.079 | 0.751 | 0.383 | 0.008 |
|  | *14370<sup>f</sup>* | *-25993.112* | *0.077* | *0.701* | *0.371* | *0.0074* |
|  | 14698 | -26114.818 | 0.076 | 0.681 | 0.370 | 0.0084 |
|  | 14847 | -26147.026 | 0.076 | 0.695 | 0.377 | 0.0086 |
|  | 15105 | -26207.243 | 0.073 | 0.634 | 0.354 | 0.0079 |
|  | 15253 | -26175.667 | 0.075 | 0.680 | 0.369 | 0.008 |
| Naïve_Opt | 13658 | -26362.214 | 0.074 | 0.61 | 0.326 | 0.01 |
|  | 13695 | -26290.942 | 0.076 | 0.662 | 0.343 | 0.0091 |
|  | 13772 | -26314.732 | 0.078 | 0.687 | 0.357 | 0.0091 |
|  | 13897 | -26324.514 | 0.079 | 0.669 | 0.345 | 0.0083 |
|  | 13912 | -26323.322 | 0.081 | 0.722 | 0.365 | 0.0084 |
|  | 13945 | -26238.085 | 0.080 | 0.699 | 0.358 | 0.0081 |
|  | 13978 | -26257.832 | 0.077 | 0.661 | 0.344 | 0.0088 |
|  | 14002 | -26279.572 | 0.079 | 0.708 | 0.363 | 0.0086 |
|  | *14010<sup>g</sup>* | *-26161.701* | *0.078* | *0.674* | *0.351* | *0.0084* |
|  | 14122 | -26336.625 | 0.076 | 0.6388 | 0.333 | 0.0088 |

**a:** Three methods for estimating the ancestral pillar order at the TGD were used: See *Methods* for details.
**b:** Number of syntenic breaks in the estimated order (see *Methods*).
**c:** ln-likelihood of the pillar data under the assumed order and the WGD- $bc^{bnf}$ model (Figure 2).
**e:** Order with the largest lnL, Global\_Break\_Opt
**f:** Order with the **overall** largest lnL across all three optimization methods.
**g:** Order with the largest lnL, Naïve\_Opt

**Supplemental Table 3:**
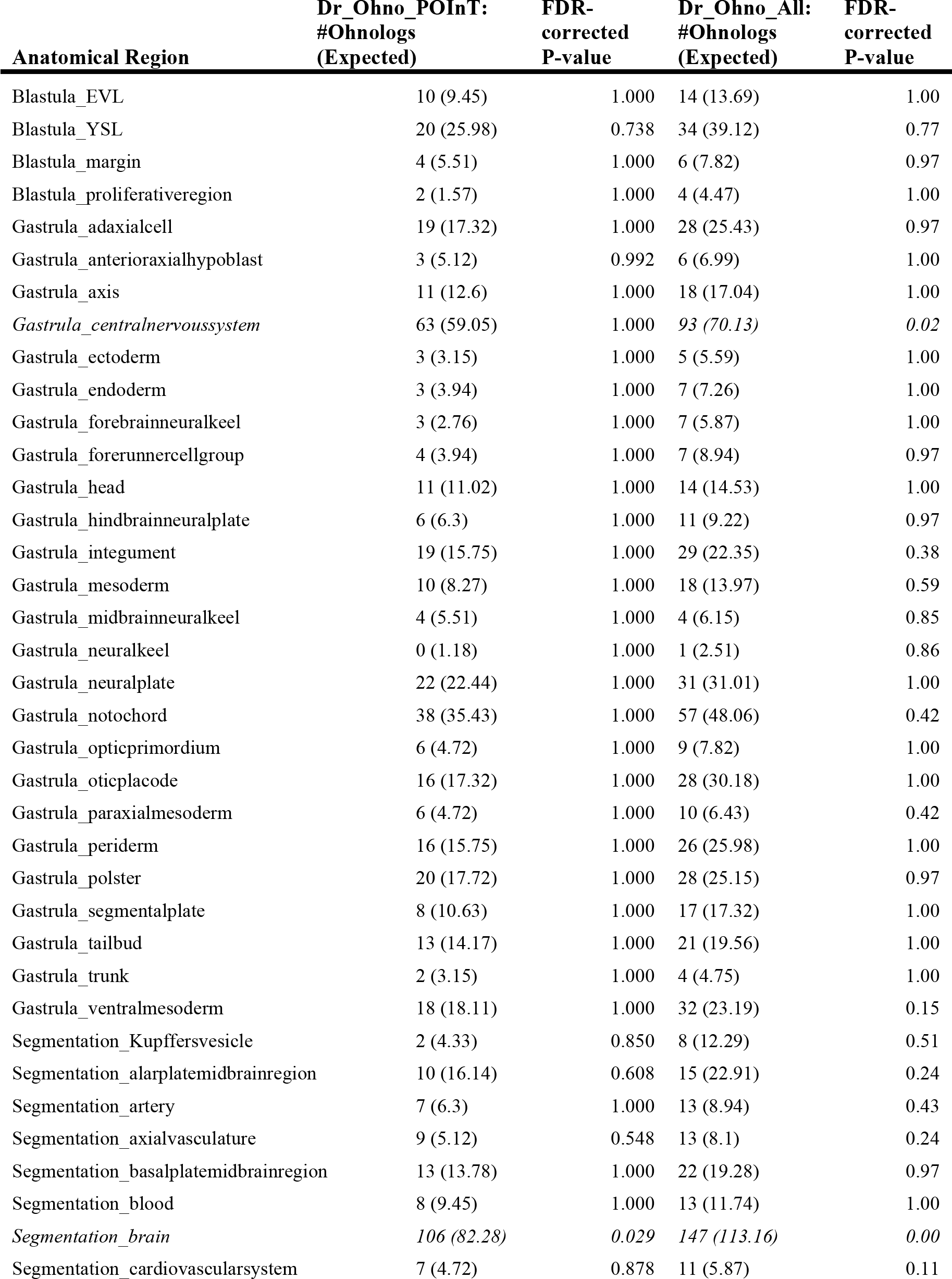
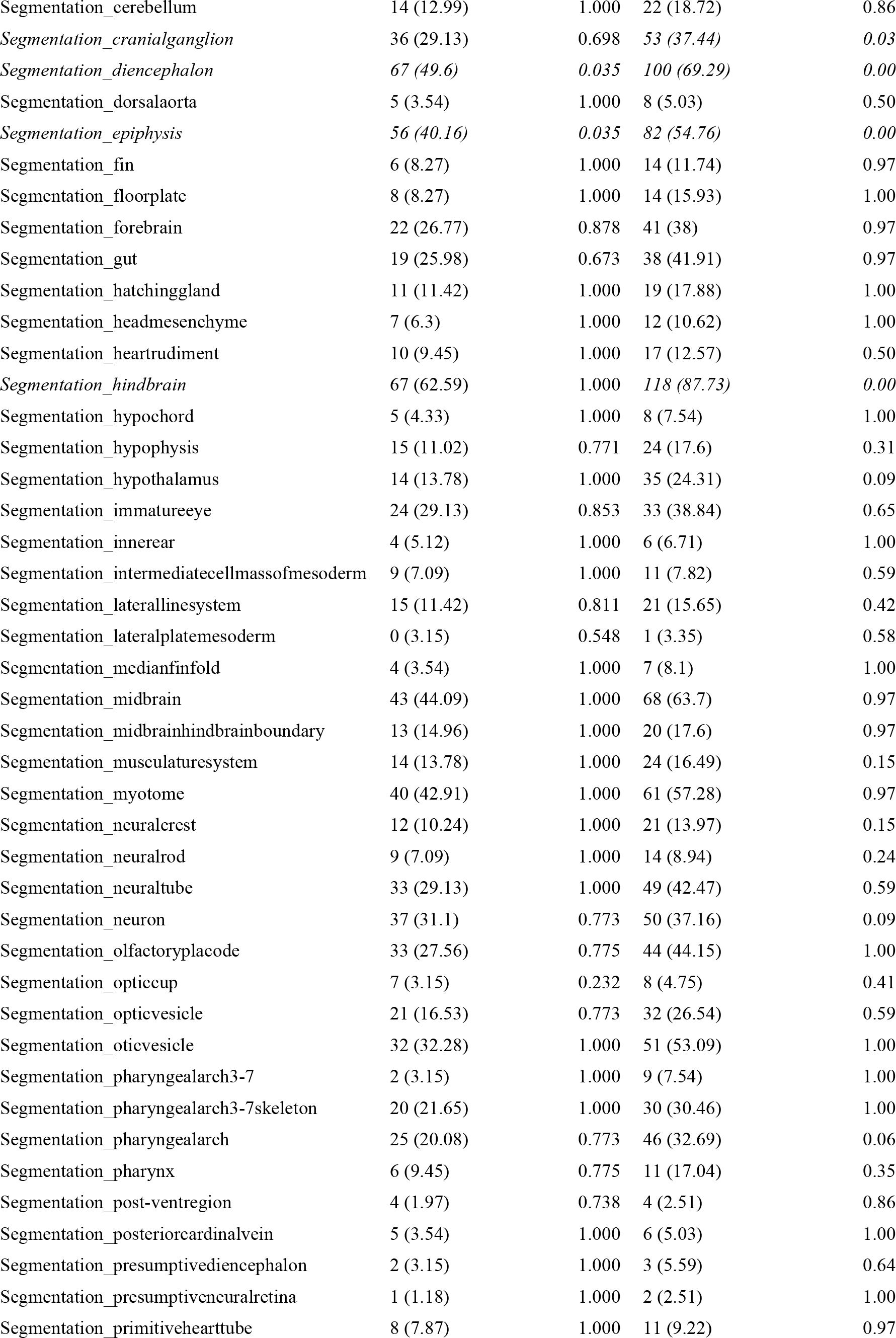
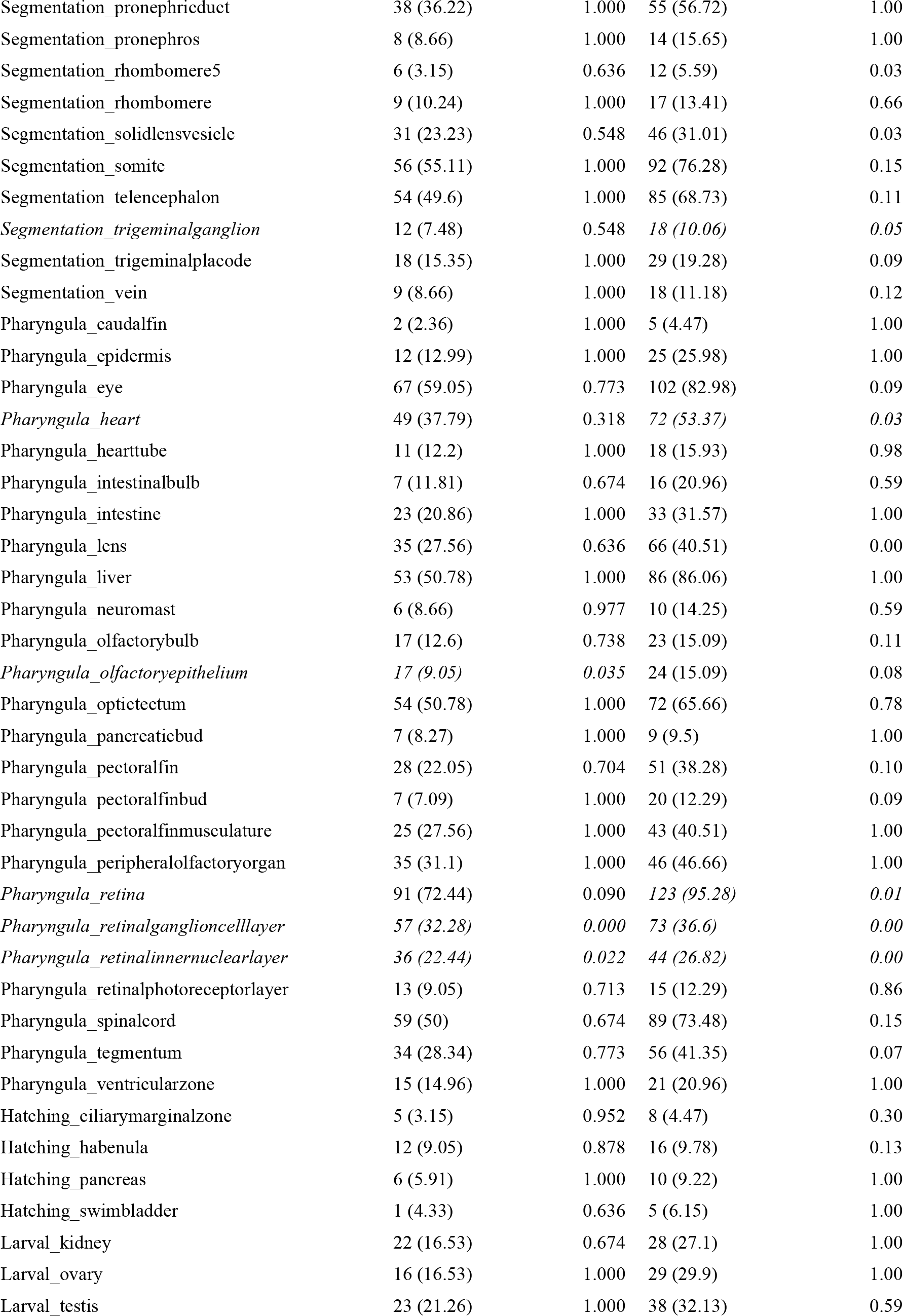
Developmental time points and anatomical regions with more or fewer ohnologs present than expected.

**Supplemental Table 4:**
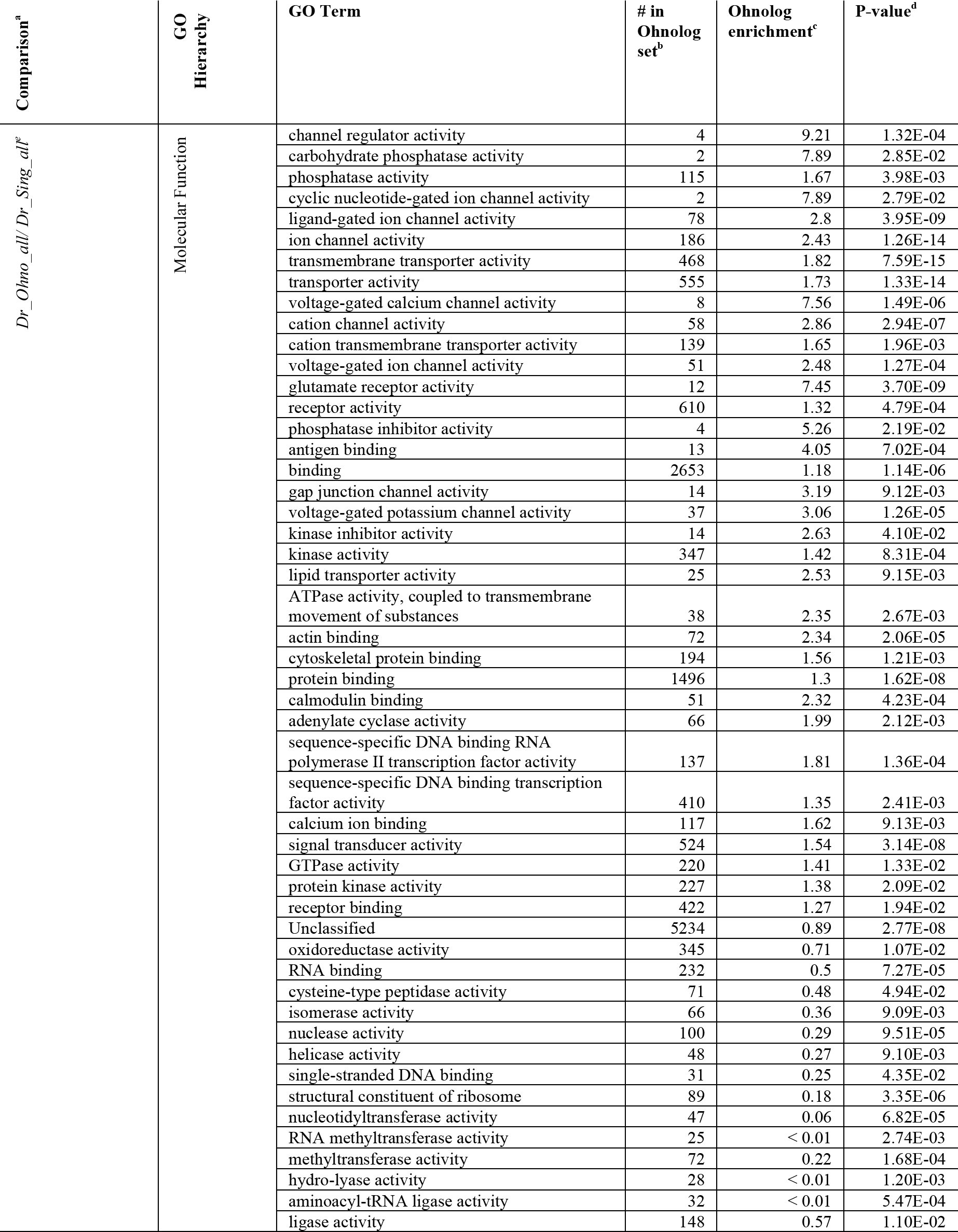
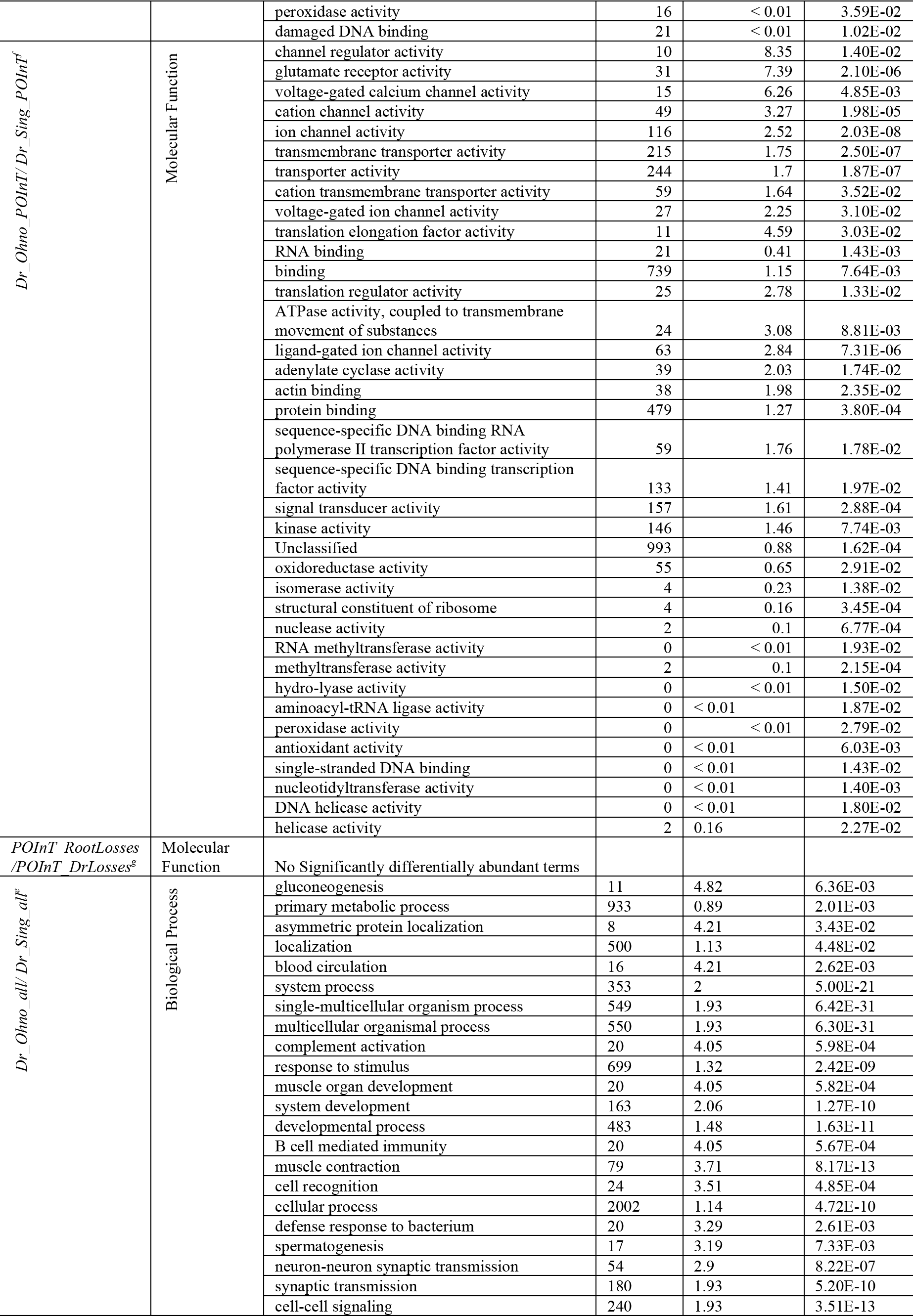
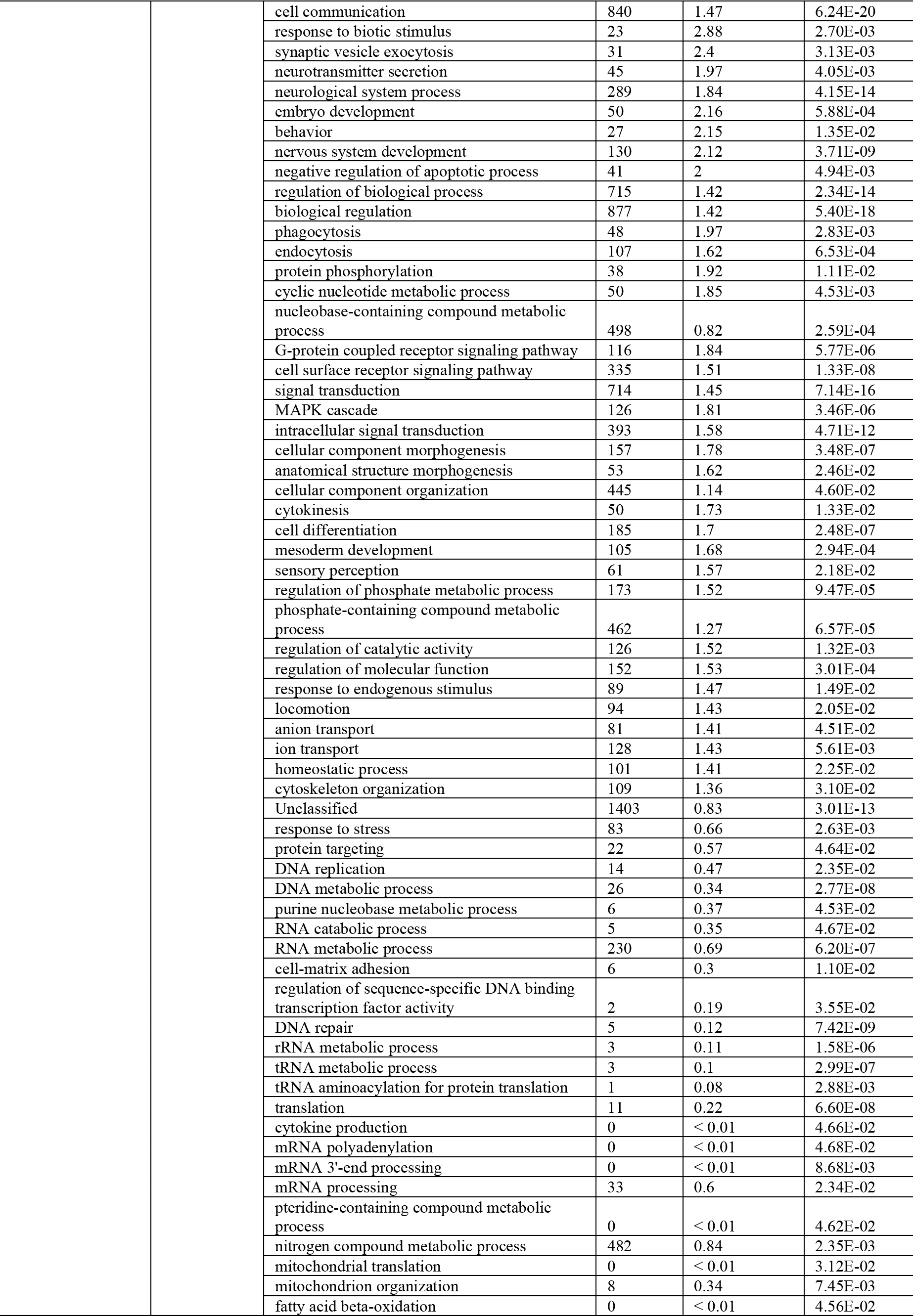
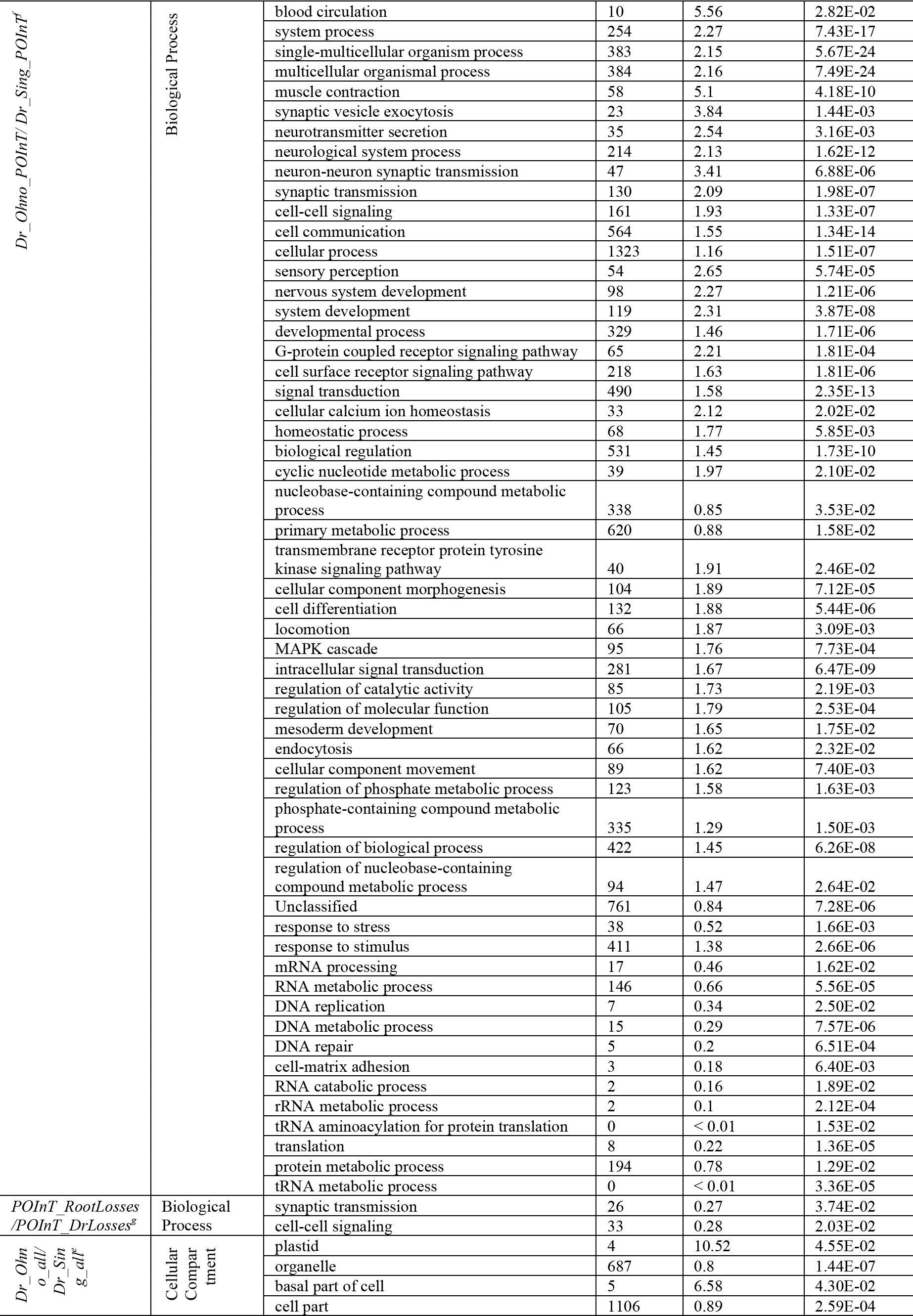
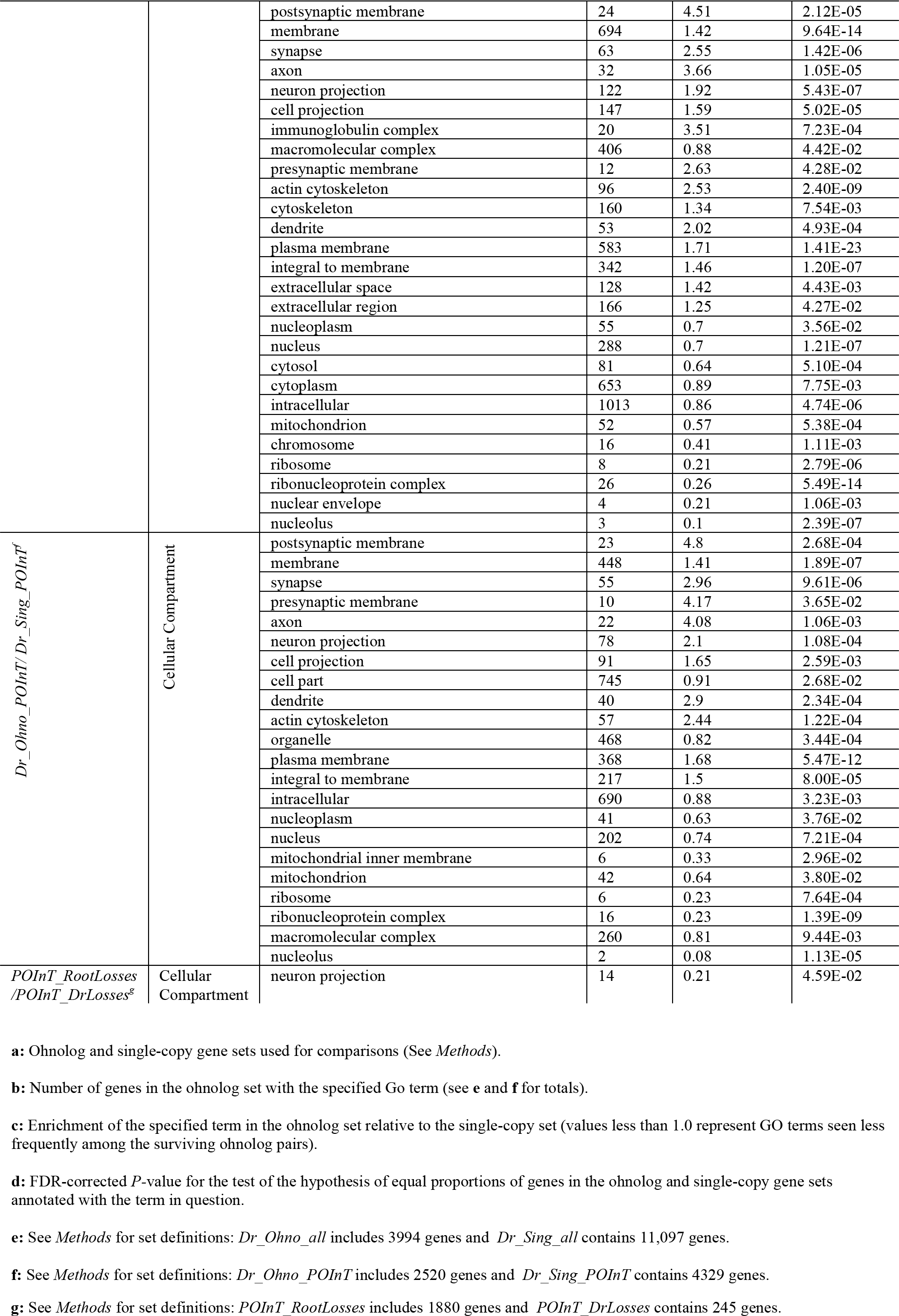
Differentially abundant GO terms for Ohnologs and single-copy genes from the TGD.

**Supplemental Data:** A compressed tar file (http://conantlab.org/data/TGD/Supplemental_data.tar.gz) consisting of nine files: 1) POInT’s orthology inferences for the 5589 pillars analyzed (tab-delimited text), giving the estimated probability of all 2^8^=256 possible orthology relationships at every pillar. 2-9) For each genome with the TGD, I give the synteny relationships seen in the estimated optimal ancestral order (tab-delimited text).

